# Social attention across development in common ravens and carrion crows

**DOI:** 10.1101/2023.08.03.551806

**Authors:** Rachael Miller, Markus Boeckle, Sophie Ridgway, James Richardson, Florian Uhl, Thomas Bugnyar, Christine Schwab

## Abstract

Underlying social learning and other important aspects of successful adaptation to social life is social awareness, where individuals are required to pay attention and respond flexibly to others in their environment. We tested the influence of social context (alone, affiliate, non-affiliate, heterospecific) on behavioural interactions (manipulation, caching, head & body out of sight i.e. barrier use) with food and objects during development at fledging (1-2 months), juvenile (3-8 months) and sub-adult (14-18 months old) in 10 carrion crows (*Corvus corone; C. cornix*) and nine common ravens (*C. corax*). These species are closely related, generalist corvids and subjects were all hand-reared and housed in highly comparable conditions. Both species will routinely cache, i.e. hide food and other items for later recovery, and engage in cache-pilfering (stealing) strategies. They will interact and ‘play’ with objects, potentially as part of developing social bonding and/or physical skills. We found that corvid behaviour was influenced by social context, with birds showing higher frequency of ‘head & body out of sight’ behaviour while others were observing than when alone, suggesting they have an awareness of other’s presence and respond by using barriers when interacting with items. There were no differences based on observer identity, supporting an interpretation of item interaction and play potentially driving development of physical skills in this setting. There were developmental effects, including increased manipulation and use of barriers as juveniles as well as increased caching with age. Ravens generally cached more than crows. Objects were manipulated more frequently than food, while barriers were used more with food, indicating that food was more likely to be actively hidden while objects may promote open play. We discuss our findings in relation to other social and developmental influences on behaviour and the wider ramifications for identifying the drivers of play in animals.

## INTRODUCTION

Social learning – learning from others - requires social awareness, where individuals recognise and respond to conspecifics and other animals within their environments, influencing who they act, react or interact with (Scheid, 2007). This ability requires paying attention to the behaviour of others and modifying their own behaviour accordingly (Heyes and Galef, 1996; Laland, 2004; Whiten et al., 1999; Whiten and Schaik, 2007). A baseline to study social interactions is through social attention: ‘a cognitive process that underlies gazing at or with another’ (Richardson and Gobel, 2015). Social attention, including monitoring others, can facilitate social learning, competition, cooperation and communication (Scheid, et al., 2007; Schwab, et al., 2008; Range, et al., 2009; Kulahci, et al., 2016). To effectively carry this out, an individual must decide where to look and how long to invest in this behaviour as it induces costs, i.e. time and habitat constraints (Klein, et al., 2009). Therefore, individuals need to select other individuals to focus on in order to optimise information-gaining processes (Wascher, et al., 2014). Their selectivity may depend on social relationships (e.g. affiliative/ non-affiliative), other aspects of identity like age, as well as the species’ social structure and life history traits (Gajdon and Stauffacher, 2001; Moscovice and Snowdon, 2006; Scheid, et al., 2007; Schwab, et al., 2008; Kulahci, et al., 2016).

Social attention has been explored quite extensively in non-human primates (e.g. Klein, et al., 2009; Schino and Sciarretta, 2016; Kano and Call, 2017; Kano, et al., 2018; Bethell, et al., 2019, Grampp, et al., 2019). For example, mandrills *(Mandrillus sphinx)* and juvenile vervet monkeys *(Chlorocebus pygerythrus)* increased their gaze on high-ranking mates compared to low-ranking mates and gaze more at their own kin than non-kin, which was also shown to relate to age (Schino and Sciarretta, 2016, Gramp, et al., 2019). Kano and Call (2017) found a variety of similarities in expression of social attention between humans and apes, adjusting their behaviours in relation to social contexts.

While social attention has also been studied in some bird species, including corvids, the role of development and focus on the focal behaviour, rather than observer behaviour, is less explored. It is of interest to explore the extent to which corvids share fundamental mechanisms of social attention with non-human primates (Emery, 2006). Carrion crows *(Corvus corone*) and common ravens *(Corvus corax)* live in flexible social groups. Social living allows animals to gain valuable social information such as; predators, food sources and mate quality (Wascher, et al., 2013). A fission-fusion social dynamic can pose some unpredictability and high variability within an individual’s social environment, but it opens up further social opportunities (Silk, et al., 2014). Ravens have one of the longest periods of socio-cognitive development of any other avian species - they are not typically successful in reproduction until their fifth year and may delay it to their 10^th^ year (Fransson, et al., 2010, Fraser and Bugnyar, 2010). Avian social systems outside of the breeding system need to be better understood as there is relatively little data on social relationships before sexual maturity or outside of the breeding season (Boucherie, et al., 2019).

The context in which social attention has been studied most often in corvids is caching – hiding food or objects for later use. Food caching may be a strategy to secure food from conspecifics as well as to save surplus food for later use (Bugnyar, 2013; Carrascal and Moreno, 1993; Vander Wall and Smith, 1987), as indicated in pinyon jays while caching in groups (*Gymnorhinus cyanocephalus)* (Marzluff and Balda, 2010). Awareness of observers and attention to their behaviour may be advantageous when competing for food and caching, which would allow flexibility in access to food and cache protection strategies (Bugnyar and Kotrschal, 2002). When manipulating the visibility of food in visual competition tasks, corvids react to what conspecifics can and cannot see (Emery and Clayton, 2002; Dally, Clayton and Emery, 2006; Bugnyar, 2011; Bugnyar et al., 2016). Gaining experience in observing others caching has been linked to successful pilfering, due to demonstrated longevity in memory of observed caches (Heinrich and Pepper, 1998) and improvement in object permanence (Bugnayr et al., 2007a).

Pilfering imposes high costs to cachers as the benefit of storing food is diminished if it gets stolen before it is retrieved (Andresson and Krebs, 1978). By engaging in countertactics, this can reduce the risk of being observed e.g. increasing distances from observers and using obstacles to restrict viewing (Bugnyar and Kotrschal, 2002). To effectively protect caches, paying attention to others is therefore vital. Pilferers have also shown to engage in displacement behaviours like digging or manipulating objects, which may function to lead others away from caches (Bugnyar and Kotrschal, 2004). As successful caching and pilfering requires complex social skills, therefore there is evidence for learning, particularly in the early stages of life (Bugnyar, Stöwe and Heinrich, 2007). For instance, for the first few months of life, up to half of a raven’s caches may contain nonedible objects, potentially to test the observer’s response and practice effective cache and cache protection in a low-risk context (Bugnyar et al., 2007a; Auersperg, et al., 2015). Indeed, when subadult ravens were confronted with unfamiliar humans that either pilfered their playfully created object caches or not, they subsequently discriminated between those humans in the food context, caching food outside view from skilled (object cache) pilferers but not from unskilled ones (Bugnyar et al., 2007b).

Social awareness is likely also a requirement during play behaviour (de Kort and Clayton, 2006; Jacobs, et al., 2014; O’Hara and Auersperg, 2017). Play allows animals to gain experience of conspecifics, their behaviour and the wider environment and is linked with increased intelligence and survival (Naples and Rothschild, 2015). Play can be sub-divided into social, locomotor and object play (Bekoff 1984; Burghardt, 2005). Locomotor play allows individuals to practice any motor-acts to improve dexterity and motor patterns, such as running or flying in the wind (O’Hara and Auersperg, 2017). Social play can aid in establishing bonds with (typically) conspecifics (O’Hara and Auersperg, 2017; von Bayern et al., 2007). Object play can allow individuals to develop skills to improve physical problem-solving, in particular, species that use tools seem to benefit from play during their ontogeny (O’Hara and Auersperg, 2017). The function of play may vary by species (Palagi, Cordoni and Tarli, 2004). For example, wild (tool-using) chimpanzees (*Pan Troglodytes*) manipulate leaves, fruit and sticks in a playful manner, which suggests this may facilitate the use of tools and aid in building nests (Pellegrini and Smith, 2005; Bjorklund and Gardiner, 2010). Corvids, such as common ravens, show complex play behaviour (Ficken, 1977). For example, object play in ravens may help young birds to learn to cache effectively in a low-risk situation, or to test social skills by encouraging pilfer interactions (Bugnyar, et al., 2007).

Play allows the development of strength, endurance and various skills through repetition of motor patterns improving an individual’s physical fitness, known as the ‘physical’ hypothesis, and through fine-tuning, this could evolve through selection as an adaptative skill to modify existing motor patterns to deal with changing situations (Bekoff, 1984). In contrast, the ‘social’ hypothesis suggests play strengthens and maintains social bonds, forming social attachments (Bekoff, 1984) (although social bonds could potentially be formed through other activities, here, we focus only on play behaviour). In wild Japanese macaques *(Macaca fuscata),* social play is important for immature subjects to build social relationships, which is fundamental for their social life (Shimada and Sueur, 2018). However, adults can play when they need to prevent or solve disputes, to anticipate or buffer forthcoming periods of social tension as well as keeping attention of a conspecific away from a resource, which serves an advantage at an immediate level or to establish good relationships in the short-term (Palagi, 2018).

The socialising function in play may be more important for some species than others (Poirier and Smith, 1974). Play within Bottlenose dolphins *(Tursiops truncatus)* and other delphinid species allows them to create novel and safer experiences for themselves and their playmates, which is seen predominantly more in calves than adults, potentially increasing the likelihood of individual survival and reproduction (Kuczaj and Eskelinen, 2014). Belding’s ground squirrel *(Spermophilus beldingi)* juveniles play almost exclusively with littermates but not at equal rates, as they tend to have one preferred partner, indicating familiarity and stability are important when regarding social play interactions (Nunes, et al., 2004). Exposure to a variety of play interactions helps facilitate motor development and individuals chose play tasks that were the greatest challenge (Nunes, et al., 2004). Social play has also been associated with larger brain mass to body mass and in relation to longer lifespans (Kaplan, 2020).

Social context and development influences exploration behaviour in ravens and crows (Miller, et al., 2015). Both species interacted most with novel items as juveniles, potentially relating to major developmental steps like dispersal, and interacted more frequently when conspecifics were present. Species also differed, for instance, in their response to a novel person, though not a familiar person (Miller, et al., 2015). On the individual level, the ravens were strongly shaped by their sibling subgroup presence, showing behavioural similarities likely driven by social context, rather than relatedness, which may facilitate social learning and cooperation (Miller, et al., 2016). These experiments focussed on responses to novelty, rather than familiar objects and food, across development. Social attention was influenced by kinship in male carrion crows, as male crows observed non-kin at a higher frequency than kin, while females showed no preference, which demonstrates that social factors affect attention patterns in carrion crows (Wascher, et al., 2014). Other studies on aspects of social attention have tended to focus on how and when individuals watch others, i.e. how they act as observers (e.g. common ravens, jackdaws; Scheid, et al., 2007; Kulahci, et al., 2016). There remain open questions regarding the behaviour of the focal, i.e. the subject who is being watched – as opposed to the behaviour of observers – i.e. whether individual ravens and crows pay attention and adjust behaviour according to who is watching them, and specifically, how this develops during ontogeny.

In this study, we presented the focal subject (raven or crow) with a familiar food or object while alone, in the presence of a sibling/ affiliate or a non-sibling/ non-affiliate conspecific, or with a heterospecific observer in the adjacent compartment. Subjects were tested in three test ‘rounds’ from fledging to sub-adult stage (1 month to ∼1.5 years old). We focused on three behavioural measures: item manipulation, item caching and – as measure of barrier use while interacting – ‘head and body out of sight’ (to observer). We aimed to investigate whether, and if so at which life stages, focal subjects altered their behaviour in the presence of a) observers generally compared with being alone and b) specific observers. By manipulating the identity and proximity (e.g. social context) of the observer and focal subject, we aimed to address the wider function of play, specifically, whether it may serve to develop physical components of competitive cache protection and/or to establish or investigate social relationships.

Furthermore, using two species that were hand-reared, housed and tested in a highly comparable manner, we were able to explore whether there were species differences in behaviour. We selected food and objects in order to assess whether subjects reacted to these items differently, assuming that food may be considered to be a higher value and therefore higher risk for stealing or pilfering by others.

Table 1 summarizes our main questions. Concerning question 1, we expected to find a difference in the focal subject’s behaviour between the alone and social contexts. If interactions contribute to the development of social relationships, we may expect subjects to behave differently depending on the identity and relationship with the observer (i.e. affiliate vs non-affiliate conspecific; conspecific vs heterospecific observer) – potentially utilising the play opportunity to learn about specific observers. Previous studies showed that having affiliates pays off in a variety of ways; increasing the individuals access to food, affects the time spent exploring new objects (Stöwe, et al., 2006), the likelihood of social learning (Schwab, Bugnyar and Kotrschal, 2008; Schwab et al., 2008) and the amount of attention spent on others (Scheid, Range and Bugnyar, 2007). Alternatively, if interactions contribute more to the development of physical skills in food competition only, i.e. to practice with item manipulation and use of barriers to prevent pilfering, we may expect an effect of social presence (compared to being alone), but no influence of different types of observers (i.e. affiliate, non-affiliate, heterospecific).

**Table 1:**
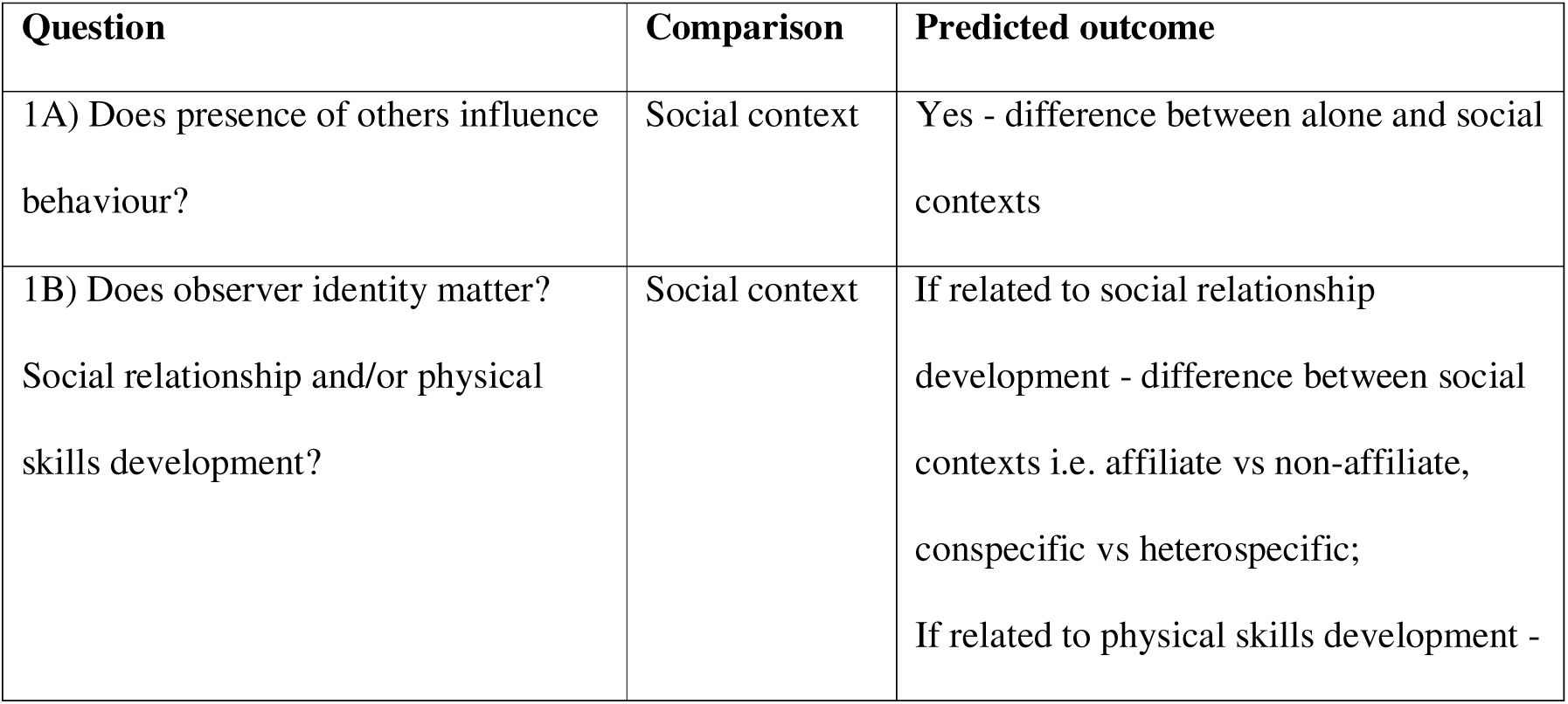
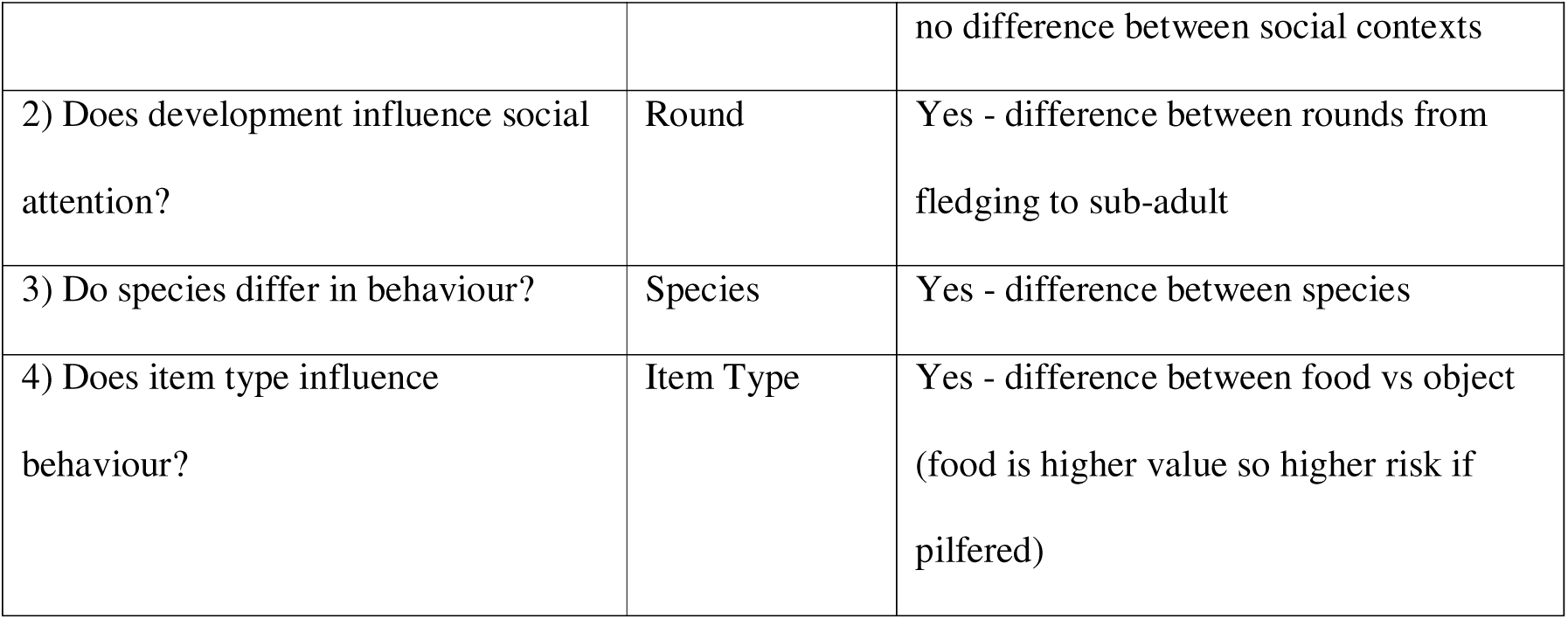
Questions, predicted outcomes and observed outcomes.

Concerning question 2 and 3, we expected developmental effects and some species differences relating to the frequency of interactions with the familiar items, similar to Miller, et al (2015)’s findings with novel items in the same corvid subjects. Furthermore, other species differences between corvids have been found. For instance, ravens were more attentive to conspecifics than jackdaws *(Corvus monedula)*, as well as showing higher interest towards food-related objects and increased duration of looks towards affiliates than non-affiliates (Scheid, Range and Bugnyar, 2007). Gallego-Abenza, Boucherie and Bugnyar (2022), noted that individuals reared in smaller family groups were more attentive than those reared in large families, suggesting upbringing can affect how juvenile ravens value social information. Therefore, we can assume that development and age could affect social attention towards ravens and potentially other corvid species, like carrion crows. For question 4, we expected objects and food may be treated differently by the focal, potentially relating to higher risk of losing food to a competitor, for instance, ravens responded to seeing others’ object manipulation, whereas jackdaws did not (Schwab et al., 2008).

## METHOD

### Subjects

Subjects were nine common ravens (six males, three females), and 10 carrion/hooded crows (four males, six females). The crows comprised of carrion/hooded crow hybrids, which reflects the overlapping range of the two subspecies from Central Europe. The ravens were obtained at pre-fledging age from three European zoos in Austria, Germany and Sweden of which five were first generation (parents wild-born) and four ravens second generation (grandparents were wild born). The crows were collected at pre-fledging age from several wild nests within Donaupark, Vienna, Austria, in 2012. Subjects were hand-reared initially in 2012 within their sibling groups until fledging and then in species groups under the same controlled conditions using the same rearers, feed, diet, training schedule, amount of human contact and social holding. The studies were comparable as much as possible for both species in the general set-up and hand-rearing.

The raven and crow group comprised three conspecific sibling groups consisting of four: three: two birds as well as one bird that did not have any siblings. They were identifiable via colour rings and were trained using positive reinforcement techniques to enable subjects to be separated within the voluntary test compartments. One raven was not included as a subject in the study as he could not be separated individually. Two crows died due to unrelated causes during the study (*Clostridia* infection, June and July 2013), so did not participate in the latter part of the study. Subjects were well habituated to people (including the experimenters: RM and CS) – important given that familiarity can positively affect the experimental performance in corvids (Cibulski, et al., 2014). Housing was organised in species groups within large, outdoor aviaries (total size ∼680m2) at Haidlhof Research Station, Austria. Individual subjects could be separated from the group in the main aviary into test compartments (∼20m2 per compartment). Visual access between compartments could be controlled by using doors and windows over the mesh.

### Procedure

There were four test conditions: individual (i.e. alone), sibling/affiliate observer, non-sibling/ non-affiliate observer and other species observer. There were three test ‘rounds’, i.e. the subjects were all tested at three separate time periods during development: Round 1 ‘fledging’ stage in June 2012, Round 2 ‘juvenile’ stage in November 2012 and Round 3 ‘sub-adult’ stage in October 2013. In Rounds 1 and 2, the observer was either a sibling or non-sibling, while in Round 3, the observer was an affiliate or non-affiliate as determined by separate social interaction focal data. The fourth condition of ‘other species observer’ involved a raven observer for the crow subjects and a crow observer for the raven subjects.

The sex of the observer was kept constant in all cases except two subjects (one female per species), as there were only male siblings. Condition order was counterbalanced within test rounds. Each test trial lasted 10 minutes, with one trial run per day over consecutive days. In Rounds 1 and 2, there were eight raven subjects and nine crow subjects in all conditions and contexts (Supplementary Table 1). In Round 3, there were nine raven subjects and eight crow subjects, although two crows died prior to this round, we could include the one single raven and one single crow that did not have siblings as the observers were affiliates/ non-affiliates rather than sibling/ non-sibling.

The focal subject was presented with two different item types: either a large piece of familiar food (Round 1: liver, Round 2: sausages, Round 3: pork) or a familiar object (Round 1: wooden blocks of neutral colour, Round 2: pinecones, Round 3: small Lego pieces). The trial began once the item had been placed on the ground in the middle of the compartment. During each trial, the focal subject was separated individually from the group into one test compartment. In the alone context trials, the focal subject was alone with the food/object with the observer out of view but present in the adjacent compartment (focal could therefore hear but not see the observer and vice versa). In the social context trials, the observer was separated individually into the adjacent compartment to the focal and visual (but not physical) access between the compartments was allowed by opening the door and window, which were covered with mesh (Figure 1). In Round 3 (sub-adult) only, the observer was first separated (five mins, with visual access only) and then was introduced into the focal’s test compartment (five mins, visual and physical access). The Round 3 set-up differed slightly from those used in Rounds 1 and 2 (where there was only visual access) to explore whether providing the observer with physical and visual access to the focal may influence focal behaviour, for instance, by increasing their perception of this as a high competition setting.

**Figure 1.**
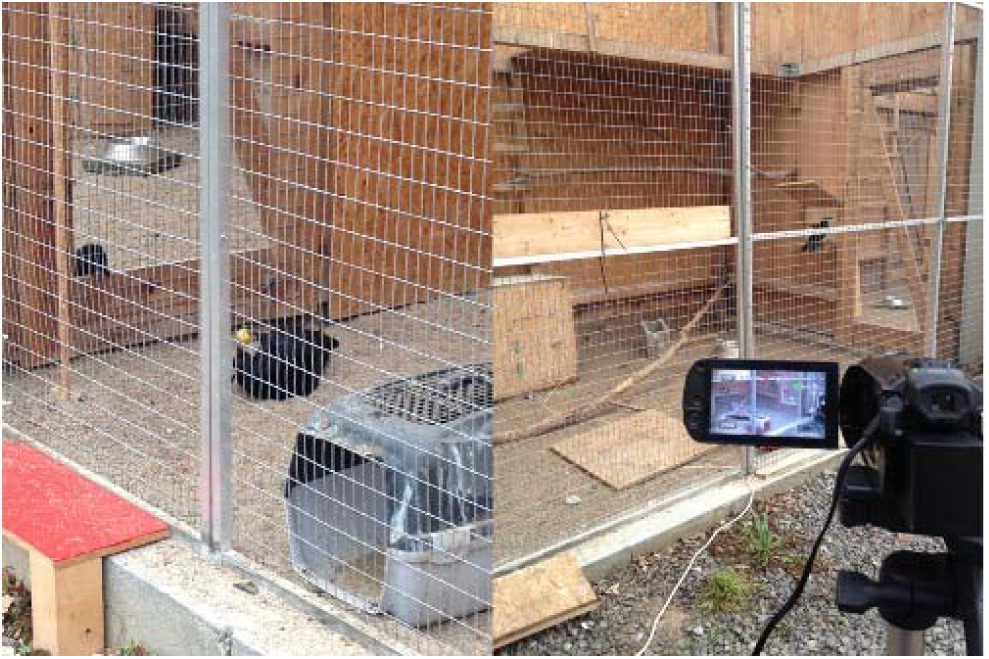
Examples of experiment set-up. Left: focal raven manipulates object in view of an observer in an adjacent compartment. Right: view of the observer and focal compartment with adjoining mesh-covered viewing door and window (multiple barriers available).

The focal test compartment was set up to provide options to cache/manipulate the item in or out of sight of the observer through the provision of visual barriers between themselves and the observer, by either occluding their entire body (e.g. a large opaque crate) or just their head (e.g. a large log) from the view of the observer (Figure 1). We recorded the frequency of item manipulation and caching behaviour of the focal subject, including their location within the compartment, as well as whether their head and body were out of sight of the observer.

Affiliation data was collected via five-minute focal observations per subject collected across all subjects, four times per week, throughout the study duration. This data included three particular measures: 1. Initiate contact sit, i.e. sitting within beak reach of another bird, 2. Allopreening, i.e. preening/ grooming the feathers of another bird 3. Touch, i.e. contact using bill/ wing/ foot with another bird. Round 3 (sub-adult) affiliation dyads were comprised using this data by pairing the focal with the observer that they showed the highest (‘affiliate’) and lowest (‘non-affiliate) frequency of affiliative behaviours (Supplementary Table 1). Round 1 (fledgling) and 2 (juvenile) dyads were siblings/ non-siblings (i.e. from the same nest or not).

### Ethical note

All procedures were conducted in accordance with the Federal Act on the Protection of Animals 2005 (Animal Protection Act TSchG). Permission to remove carrion crow nestlings from the wild was received from the Magistrate of Vienna for Environmental Protection (MA 22-355/2012.4) causing minimum disturbance and stress and in accordance with the Animal Transportation Act 2007 for both species (Crows: 10-14 days old; Ravens: 25-38 days old). Only individuals deemed healthy were removed from the nest during breeding season. Both species are listed as Least Concern (BirdLife International, 2016; 2020). Therefore, this had a minimal impact on the population and encouraged double clutching. Personnel responsible for the removing, handling and transporting of subjects were professionals and provision of ventilation and space was provided along with being hand-fed adequate food and water upon arrival. Subjects were nine common ravens (six males, three females), and 10 carrion/hooded crows (four males, six females), which were hand reared within their sibling groups to minimise stress and then in social species groups. The birds received suitably sized leg rings for identification purposes.

Habituation and positive reinforcement training were carried out immediately by trained professionals upon arrival of the subjects to minimise stress. Standard species husbandry practices were met throughout, and a variety of species-specific enrichment choices were provided to improve animal welfare, maximising welfare and survival by encouraging natural behaviours. These enrichment types included food and novel objects to encourage problem-solving, social (conspecifics, heterospecific, human interaction) and physical (branches, substrate). Individual behaviours were also assessed upon arrival and through-out the study to identify any atypical or aggression-related behaviours. Stressors were shown by one individual raven when encouraged to use rewards to be temporarily separated from the group into the test compartment; therefore, this individual was excluded from further testing.

Permission to house animals at Haidlhof Research Station was obtained from the Austrian Ministry for Science and Research (BMWFW-66.006/0011-WF/II3b/2014). This study was reviewed and approved by the Animal Welfare Board at the Faculty of Life Sciences, University of Vienna (2014.012). Subjects were housed in species groups within large, outdoor aviaries (total size ∼680m2) at Haidlhof Research Station, Austria (University of Vienna and University of Veterinary Medicine), providing suitable temperatures, photoperiod and shelter. The test compartments still allowed enough autonomy for the subjects (∼20m2 per compartment). Prior to testing, all subjects were satiated with 50% of their daily diet, consisting of a mix of fruit, meat, vegetables, bread and yogurt and water was constantly available to control for any differences in subject motivation driven by thirst or hunger. With only one trial run for this study per day (10 minutes long), with voluntary participation (i.e. birds were encouraged – but never forced - to enter the test compartment for a food reward), this caused minimal disturbance.

The number of subjects is comparable to other relevant published data (e.g. Scheid, Range and Bugnyar, 2007 – 9 birds; Bugnyar, Reber and Buckner, 2016 – 10 birds) and were chosen to test the hypothesis without the loss of scientific rigor i.e. variability of populations or invalidity. Several other non-invasive, behavioural studies were conducted on these subjects (e.g. Miller, et al., 2015; Miller, et al., 2016) allowing for past and potential future publications. All applicable Austrian and University of Vienna animal care and use guidelines were followed to ensure adequate animal welfare through-out the study. The study did not involve any invasive or potentially harmful techniques as positive reinforcement was used and the individuals only participated with free will, providing autonomy. All the subjects remained at the Haidlhof Research Station after the study was completed to allow for further behavioural studies, providing exceptional welfare through-out. Releasing the crows back into the wild would have been detrimental to the individuals due to human reinforcement training, existing populations, potential spread of disease and competition. Using subjects that were either wild-caught or first generation wild, reared in a similar manner, allows for increased value and quality of the outputs in reference to wider populations, resulting in a beneficial and invaluable study enhancing understanding of natural social behaviours of these two corvid species.

The authors declared that they have no conflicts of interest.

### Data Analysis

RM and CS collected data. Videos were coded using Solomon Coder by FU and JR, with both coders first coding a sub-set of videos with excellent inter-rater reliability (Spearman ρ > 0.991, *p*<0.001). We used General Linear Mixed Models (GLMM) with negative binomial distribution to account for the Zero-Inflation. To analyse the data we used all 3 rounds, with round (1 = fledgling, 2 = juvenile, 3 = sub-adult), item type (0 = object, 1 = food), social contexts (1 = alone, 2 = sibling/affiliation, 3 = non-sibling/non-affiliate, 4 = heterospecific) and species (0 = common raven, 1 = carrion crow) as main effects, species*social context and species*round as interaction effects, and focal ID as a random effect. We focused on three behavioural measures of interest: frequency of item manipulation, item caching and head & body out of sight (latter as a measure of barrier use during item interaction/ caching). For each behavioural measure we ran 2 GLMMs; first the full model as presented above and a model with the random effect alone. We used likelihood ratio tests to compare the full model (all predictor variables, random effects) with the null model (random effects only). We used Tukey comparisons for post-hoc comparisons of the full model.

Statistical analyses were carried out using R 4.3.1 (R Core Team, 2023). Zero-inflated GLMMs were calculated with the glmmTMB package (Brooks, et al., 2017); posthoc comparisons were calculated with emmeans (Lenth, et al., 2023); to check distribution and residual diagnostics the package DHARMa (Hartig and Lohse, 2022) was used.

### Data Availability

The full data sheet and R script are available via Figshare: DOI 10.6084/m9.figshare.23642847 (private link: https://figshare.com/s/ed4275e740d22444af35).

## RESULTS

Item manipulation differed between item type, species*round and species*context, though not context, species or round (GLMM Supplementary Table 2) and the full model differed from the null model (*X*^2^ = 74.302, df=12, p<0.0001). Caching differed between species, round, and species* round with a non-significant trend for context, item type and species*context (GLMM Supplementary Table 2) and the full model differed from the null model (*X*^2^ = 111.14, df=12, p<0.0001). Head and body out of sight differed by context, item type, round, species*round, though not species or species*context (GLMM Supplementary Table 2) and the full model differed from the null model (*X*^2^ = 121.11, df=12, p<0.0001). A summary of pairwise comparisons is outlined in Table 2 (full output in Supplementary Table 3 and 4).

**Table 2.** Summary of effects per behaviour measure following Tukey pairwise comparisons. * = p<0.05. ns = p>0.05. R=raven, C= crow.

| Effect/<br>Behaviour | Item Manipulation |  | Item Caching |  | Head & Body Out of<br>Sight |  |
| --- | --- | --- | --- | --- | --- | --- |
| Item Type | * | Object > food | ns | (Trend) | * | Object < food |
| Social Context | ns | / | ns | (Full model only) | * | Alone < affiliate;<br>alone < non-affiliate;<br>alone < other<br>species |
| Species | ns | / | * | Raven > crow | ns | / |
| Round | * | Fledging < juvenile;<br>juvenile > sub-adult | * | Fledging < juvenile; fledging < sub-adult;<br>juvenile < sub-adult | * | Fledging < juvenile; fledging > sub-adult; juvenile > sub-adult |
| Species*Social Context | ns | (*Full model only*) | * | Raven affiliate > crow<br>heterospecific | * | (*Trend*) |
| Species*Round | * | **Raven:** ns<br>**Crow:** fledging < juvenile;<br>juvenile > sub-adult<br>**Between species:** fledging R < juvenile C;<br>juvenile C > sub-adult R | * | **Raven:** fledging < sub-adult; juvenile < sub-adult<br>**Crow:** fledging < juvenile; fledging < sub-adult<br>**Between species:** fledging R < fledging C;<br>fledging C < juvenile R;<br>fledging C < sub-adult R; juvenile R < sub-adult C; sub-adult R > sub-adult C | * | **Raven:** fledging < juvenile; juvenile > sub-adult<br>**Crow:** fledging < juvenile; fledging < sub-adult, juvenile > sub-adult<br>**Between species:** fledging C < juvenile R; fledging C < sub-adult R, juvenile R < sub-adult C |

### Influence of social context

For head & body out of sight, there was a higher frequency in affiliate than alone (Tukey contrasts: z=-3.771, p=0.0009), non-affiliate than alone (z=-3.662, p=0.0014) and heterospecific than alone context (z=-3.012, p=0.0138) (Figure 2). Within species, for caching, there were higher caching frequencies for raven affiliate than crow other species context (z=3.656, p=0.0063).

**Figure 2.**
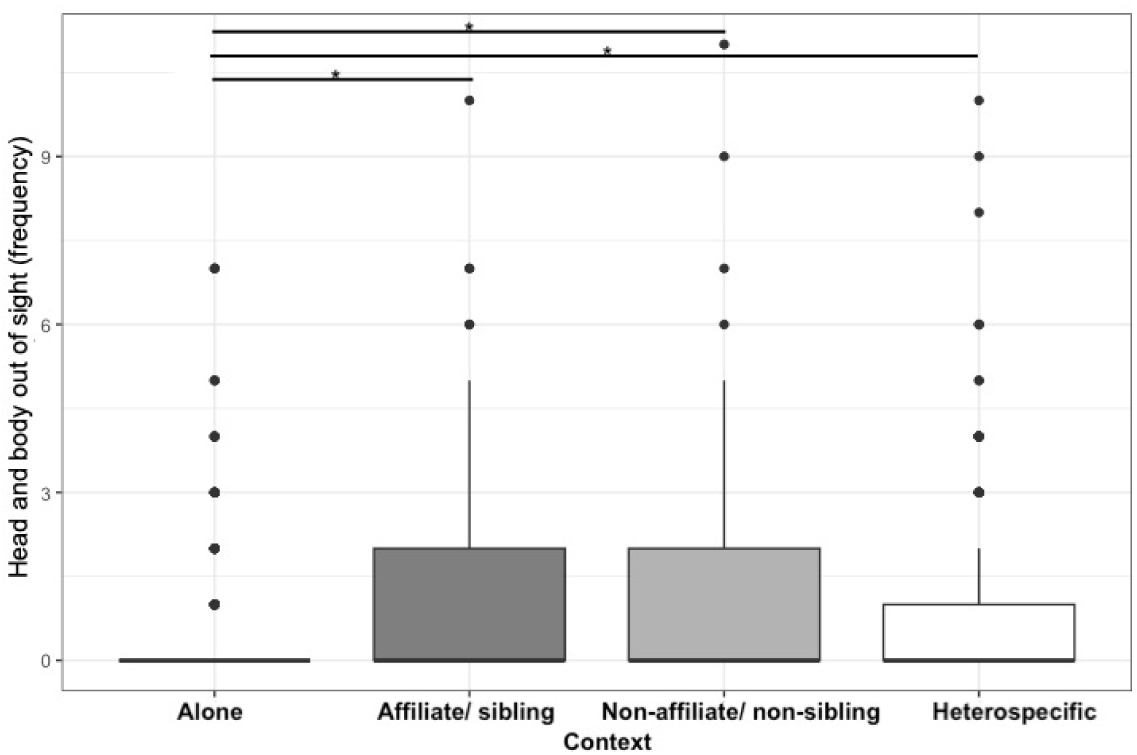
Influence of social context on head and body out of sight behaviour measure.

### Influence of round

There were effects of round for all 3 behaviour measures. There were higher frequencies of manipulation at juvenile than fledging stage (Tukey contrasts: z=-4.185, p=0.0001), and in juvenile than sub-adults (z=3.229, p=0.0036). Caching was higher in sub-adult than juvenile (z=-3.980, p=0.0002), sub-adult than fledging (z=-8.706, p<0.0001) and juvenile than fledging (z=-5.816, p<0.0001) (Figure 3). Head and body out of sight occurred more frequently at juvenile than fledging stage (z=-7.258, p=<0.0001), juvenile than sub-adult (z=4.649, p=<0.0001), and fledging than sub-adult (z=-2.613, p=0.0244). Within species, there were also multiple effects of round across all 3 measures (Supplementary Table 4). For example, there was higher caching for ravens than crows as fledglings (z=3.908, p=0.0013) and as sub-adults (z=2.984, p=0.0339).

**Figure 3.**
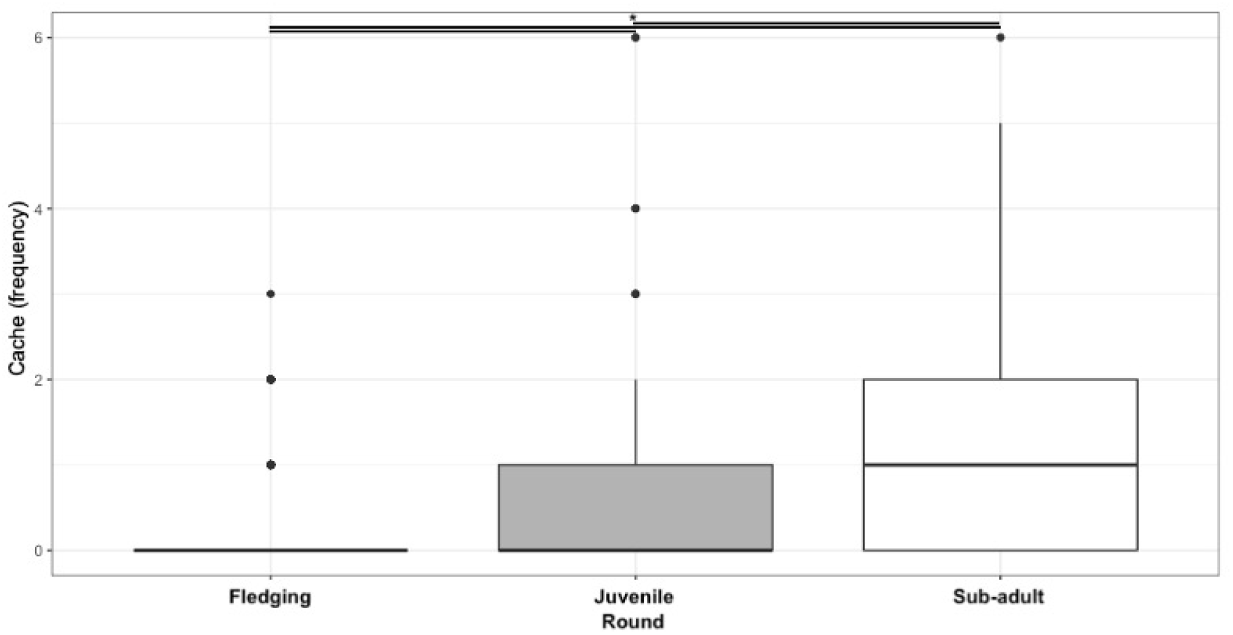
Influence of round on caching behaviour measure. Round 1 = fledging; 2 = juvenile; 3 = sub-adult.

### Influence of species and item type

Between species, ravens cached more frequently than crows (Tukey contrasts: z=3.301, p=0.001). There were effects of item type with higher manipulation frequency with object than food (z=6.516, p<0.0001; Figure 4), and higher head and body out of sight frequency with food than objects (z=-7.468, p=<0.0001).

**Figure 4.**
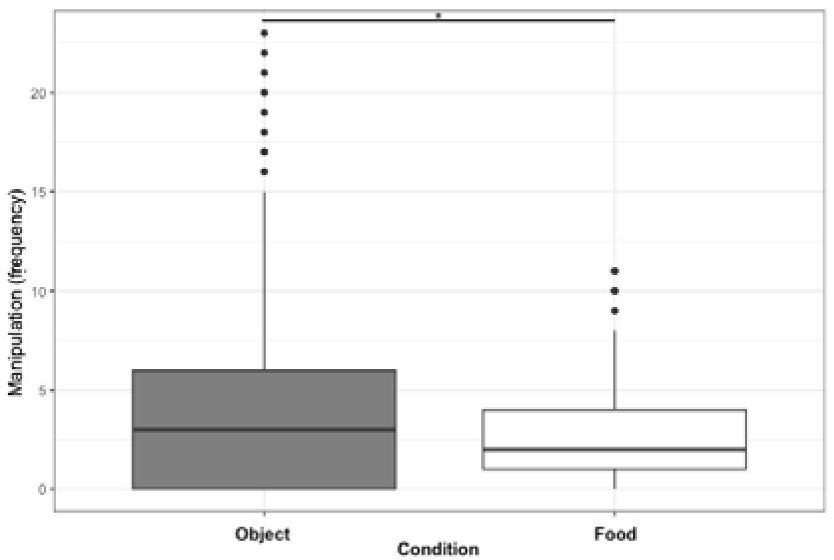
Influence of item type on manipulation behaviour measure.

## DISCUSSION

In this study, we investigated social attention across development in common ravens and carrion crows. We examined the influence of 1) social context - observer presence and identity (kin/ affiliation/ species); 2) development (fledging, juvenile, sub-adult); 3) species; 4) item type (food, object) on caching, item manipulation and head & body out of sight (i.e. use of barriers) behaviours. Our findings generally supported our predictions (Table 1). Specifically, we found: 1) support that presence of others influences behaviours, with a higher frequency of head & body out of sight in all social contexts than when alone, though no differences in behaviour based on type of observer present. 2) Developmental influences with differences in behaviour across rounds from fledging to juvenile to sub-adult for frequency of manipulation, item caching and head & body out of sight measures. 3) Species differences in caching with ravens caching more frequently than crows. 4) Item type effects with higher manipulation frequency with objects than food and higher head & body out of sight frequency with food than objects.

In relation to social context, the presence of others (any observer) led to increased use of barriers while interacting with food and objects. These findings are in line with previous studies in ravens, for instance, ravens have shown to be i) highly sensitive when in company of conspecifics who may pilfer caches (Heinrich and Pepper, 1998), ii) to reduce the time they cache and use obstacles as visual barriers when potential observers are in the vicinity (Bugnyar and Kotrschal, 2002) and iii) to consider the visual perspective of others for adjusting their caching behaviour (Bugnyar, Reber and Buckner, 2016).

These findings suggest that the birds used these ‘play’ opportunities, i.e. to manipulate objects and food and practice barrier use in presence of others, for physical skills development, rather than necessarily social relationship development. Rather than choose to engage in social play or bonding, such as interacting with items in view of observers, the birds tended to utilise barriers to hide their head and body while interacting, compared to when they were alone. As successful caching and pilfering requires complex social and physical skills, therefore there is strong evidence for learning, especially in the early stages of life (Bugnyar, Stöwe and Heinrich, 2007). Previous studies have shown that scrub jays (*Aphelocoma californica*) and willow tits (*Poecile montanus)* will adjust their frequency of caching when social competition is high (Emery and Clayton, 2002; Lahti, et al., 1998). It has been previously suggested that social attention influences kinship in male carrion crows (Wascher, et al., 2014), which was not observed in the present study. Typically, carrion crows form non-breeding flocks up to adulthood and then separate into breeding pairs (Clayton and Emery, 2007). However, in Spain, this population of carrion crows tested in Wascher, et al. (2014) form co-operative breeding groups of up to nine individuals, which consist of the breeding pair, offspring and immigrant males (Baglione, et al., 2003). Thereby there were differences in social structures between the carrion crow samples, which may require males extracting more valuable social information from non-kin (Wascher, et al., 2014). Horn et al., (2020) also found different patterns in prosocial behaviour between eight cooperative, colonial and territorial corvid species, as well as some sex effects, including stronger propensity for prosociality in males (though not female) colonial nesting species. Scheid, et al., (2007) and Schwab, Bugnyar and Kotrschal, (2008) showed that ravens are more attentive to affiliated individuals compared to non-affiliated by increasing duration rather than frequency of gazes, however, kinship and affiliation status did not affect attention in jackdaws.

A non-affiliate/ non-kin may be perceived as higher competition; however, we did not find differences in caching (or other behaviours) between affiliate/kin and non-affiliates/non-kin. It is possible that – as the observer was largely located in an adjacent rather than the same compartment – that the focal did not perceive this as a high competition setting. We adjusted the protocol in round 3 (sub-adult) to allow the observer access into the focal compartment for the second half of the trial to explore this as an explanation. There was a higher frequency of caching in round 3 than round 2 (juvenile) or 1 (fledgling), however, it is difficult to differentiate potential development effects from the procedure change. In the wild, juvenile ravens have shown to be less likely to cache then subadults or adults, and wild juveniles also have less access to food than adults, so caching opportunities may reflect food accessibility (Beck, Loretto and Bugnyar, 2020). Such findings suggest our results may relate rather to developmental changes.

We found developmental influences on behaviour with familiar objects and food, in line with previous studies in corvids focussing on responses to novel objects (Miller, et al., 2015; 2016). For instance, increased head and body out of sight behaviour at the juvenile stage (round 2) in both species correlates with an increase in use of barriers during caching juvenile ravens (Bugnyar, et al., 2007). Bugnyar, et al (2007) noted that it takes ravens a long time to show full flexibility in their ‘deception’ manoeuvres while caching. However, concealing objects (hiding from view) came almost at the same age in all subjects, occurring around the same time that ravens can succeed in geometrical gaze following around barriers (Schloegl et al., 2007). This could potentially indicate that it is a development step (Bugnyar, Stöwe and Heinrich, 2007) and could allow for controlling intentions (Bugnyar, 2013). It takes ravens three to four years to reach sexual maturity and crows two to three years, however, studies have shown independence and dispersal occurs at similar points within the same life stages (Cramp and Perrins, 1994; Ratcliff, 1997), therefore the life stages selected were comparable between species.

With regard to species, only caching appeared to differ between species, whereas frequency of item manipulation and barrier use were similar in both species. Subjects were reared, housed and tested in a highly comparable manner, though they did differ in source, with crows being wild-born and the ravens being first-generation captive (parents wild-born). Both species are closely related, with similar socio-ecology, including diet, range, large flock sizes and social structures (fission-fusion dynamics: Uhl et al., 2019; Loretto, et al., 2017). They also show similar levels of neophobia (responses to novelty) (Miller, et al., 2022). Corvid species do range in caching propensity from moderate to specialised, with the common ancestor of all corvids likely being a moderate cacher (de Kort and Clayton, 2006). Both carrion crow and common raven were categorised as moderate cachers (de Kort and Clayton, 2006), although this is one of the first studies to directly compare caching in both species using the same set-up in similarly reared and housed subjects.

A previous study indicating species differences in social attention in ravens and jackdaws found that ravens were slower in their movements and behaviours, spending more time observing and having an increased attention span (Scheid at al., 2007). This has been suggested as being potentially due to their size difference as ravens are three times larger than jackdaws (Heinroth & Heinroth, 1926). However, there are also size differences in ravens (wingspan: 120-150cm; weight 800-1500g) and carrion crows (wingspan: 93-104cm; weight 370-650g; RSPB, 2023a; 2023b), yet they appeared similarly influenced by social context.

Food and objects evoked different behavioural responses with objects being manipulated more frequently and food being more likely to lead to barrier use. These findings may relate to the risk level or value associated with each. Objects may be played with to help young birds learn to cache and pilfer effectively in a low-risk setting to develop cache-protection strategies (Bugnyar, et al., 2007). For instance, caching and re-caching in different locations or manipulating objects in view of observers. On the other hand, food may be hidden well using a barrier to hide the body and head of the bird and not re-covered/ re-cached to reduce chances of being pilfered. In ravens, half of all caches in the first few months of life contain non-edible objects, though they continue to use objects for caching and pilfering into adulthood (Bugnyar, et al., 2007).

The observer was free to move around their large adjacent compartment, so at times, they may have been away from the mesh-covered window and door (though viewable to the focal). It is possible that the focal did not therefore perceive the observer as a potential competitor in these cases. Ravens will guard caches only when a peephole is open but not when it is closed, which suggests they can generalise their own perceptual experience to conclude the possibility of being observed by others (Bugnyar, Reber and Buckner, 2016). However, this should not influence observer identity. We changed the observer from kin/non-kin in round 1 (fledging) and 2 (juvenile) to affiliate/non-affiliate observer in round 3 (sub-adult) to follow the likely changes in social structure at this stage. Corvid juveniles affiliate with multiple individuals of the same and/or opposite sex, often favouring siblings. However, as they reach sub-adult, they will spend more time with non-kin individuals of the opposite sex, which eventually results in a bonded pair (de Kort, Emery and Clayton, 2006; Scheid, Schmidt and Noë, 2008; Loretto, Fraser and Bugnyar, 2012).

Furthermore, a limitation of the study - also applicable to some other social attention studies in birds - is that birds have an aspherical view (Fite and Rosenfield-Wessels, 1975), therefore, this makes it challenging to determine their gaze (Dukas and Kamil, 2000). For example, earlier studies assumed that ground-feeding birds could not detect predators whilst in the head-down position, though this was later disproved (Lima and Bednekoff, 1999).

However, when in the head-down position, they may struggle to detect predators quickly enough in order to escape (Dukas and Kamil, 2000). Future work may aim to quantify observer attention or measure gaze direction.

It is possible that selecting an agonistic observer (highest number of negative interactions) – rather than a non-affiliate (least number of positive interactions) may illicit an identity specific response – an avenue for future research. Another avenue for future research is to focus on individual differences in social attention, as for example, those that are more social may be more efficient at gaining information compared to more aggressive ones (Taborsky and Oliveira, 2012).

It is still not clear whether these behavioural patterns persist similarly later on in life, therefore, follow up studies may compare social attention in both species as adults, although it may be more difficult to mix non-affiliates since they are typically territorial and housed in breeding pairs. Additionally, future comparison with other corvid species differing in socio-ecology (e.g. more or less social species). In humans, Oliva et al., (2003) determined that there were two different types of influences on social attention; top-down - views on other cognitive states i.e. aspirations, intentions and desires, and bottom-up - presence of another i.e. gaze direction, social identity. We have focussed rather on the latter, though future work may expand on top-down influences on social attention in other species.

In conclusion, we found influences of social context (social presence), development, species and item type (food, object) on behaviour in two corvid species. The presence of (any) observer influences behaviour from fledging. These findings expand on our understanding of social attention, developmental changes and potential drivers of (food and object base) play behaviour.

## ACKNOWLEDGEMENTS

Thanks to Martina Schiestl for animal training input and the Haidlhof Research Station staff for animal care. R.M. was supported by the Vienna Science and Technology Fund (WWTF) through project CS11-008 to C.S. and T.B. (2012-2015) and the Austrian Science Fund (FWF) through project Y366-B17 to T.B. S.R. was supported by QR funds at Anglia Ruskin University awarded to R.M (2023).

## AUTHOR CONTRIBUTIONS

R.M., T.B. and C.S. conceived and designed the experiments. R.M. and C.S. collected the data and F.U. and J.D. coded the videos. R.M. and M.B. planned the analysis and M.B. analysed the data. R.M. interpreted the data and R.M and S.R. prepared the figures and tables. R.M. and S.R. wrote the first draft of the manuscript, with subsequent drafts being reviewed by the other authors. R.M, T.B. and C.S. provided direct funding support. All authors gave final approval for publication and agreed to be accountable for all aspects of the work.

## Notes

https://dx.doi.org/10.6084/m9.figshare.23642847

https://figshare.com/s/ed4275e740d22444af35

### Competing Interest Statement

The authors have declared no competing interest.

## REFERENCES

Andersson, M. and Krebs, J., 1978. On the evolution of hoarding behaviour. Animal Behaviour, [online] 26, pp.707–711. https://doi.org/10.1016/0003-3472(78)90137-9.

Auersperg, A.M.I., Van Horik, J.O., Bugnyar, T., Kacelnik, A., Emery, N.J. and Von Bayern, A.M.P., 2015. Combinatory actions during object play in psittaciformes *(Diopsittaca nobilis, Pionites melanocephala, Cacatua goffini)* and corvids *(Corvus corax, C. monedula, C. moneduloides)*. *Journal of Comparative Psychology*, [online] 129(1), pp.62–71. https://doi.org/10.1037/a0038314.

Baglione, V., Canestrari, D., Marcos, J.M. and Ekman, J., 2003. Kin Selection in Cooperative Alliances of Carrion Crows. *Science*, [online] 300(5627), pp.1947–1949. https://doi.org/10.1126/science.1082429.

Beck, K.B., Loretto, M.-C. and Bugnyar, T., 2020. Effects of site fidelity, group size and age on food-caching behaviour of common ravens, Corvus corax. *Animal Behaviour*, [online] 164, pp.51–64. https://doi.org/10.1016/j.anbehav.2020.03.015.

Bekoff, M., 1984. Social Play Behavior. *BioScience*, [online] 34(4), pp.228–233. https://doi.org/10.2307/1309460.

Bethell, E., Kemp, C., Thatcher, H., Schroeder, J., Arbuckle, K., Farningham, D., Witham, C., Holmes, A., MacLarnon, A. and Semple, S., 2019. Heritability and maternal effects on social attention during an attention bias task in a non-human primate, Macaca mulatta. [preprint] Medicine and Health Sciences. https://doi.org/10.32942/OSF.IO/5NZD4.

BirdLife International, 2016. IUCN Red List of Threatened Species: Corvus corone. IUCN Red List of Threatened Species. [online] Available at: <https://www.iucnredlist.org/en> [Accessed 20 June 2023].

BirdLife International, 2020. IUCN Red List of Threatened Species: Corvus corax. IUCN Red List of Threatened Species. [online] Available at: <https://www.iucnredlist.org/en> [Accessed 20 June 2023].

Bjorklund, D.F. and Gardiner, A.K., 2010. Object Play and Tool Use: Developmental and Evolutionary Perspectives. [online] Oxford University Press. https://doi.org/10.1093/oxfordhb/9780195393002.013.0013.

Boucherie, P.H., Loretto, M.-C., Massen, J.J.M. and Bugnyar, T., 2019. What constitutes “social complexity” and “social intelligence” in birds? Lessons from ravens. *Behavioral Ecology and Sociobiology*, [online] 73(1), p.12. https://doi.org/10.1007/s00265-018-2607-2.

Brooks, M., E., Kristensen, K., Benthem, K., J.,van, Magnusson, A., Berg, C., W., Nielsen, A., Skaug, H., J., Mächler, M. and Bolker, B., M., 2017. glmmTMB Balances Speed and Flexibility Among Packages for Zero-inflated Generalized Linear Mixed Modeling. *The R Journal*, [online] 9(2), p.378. https://doi.org/10.32614/RJ-2017-066.

Bugnyar, T., 2011. Knower–guesser differentiation in ravens: others’ viewpoints matter. *Proceedings of the Royal Society B: Biological Sciences*, [online] 278(1705), pp.634–640. https://doi.org/10.1098/rspb.2010.1514.

Bugnyar, T., 2013. Social cognition in ravens. *Comparative Cognition & Behavior Reviews*, [online] 8, pp.1–12. https://doi.org/10.3819/ccbr.2013.80001.

Bugnyar, T. and Heinrich, B., 2005. Ravens, *Corvus corax*, differentiate between knowledgeable and ignorant competitors. *Proceedings of the Royal Society B: Biological Sciences*, [online] 272(1573), pp.1641–1646. https://doi.org/10.1098/rspb.2005.3144.

Bugnyar, T. and Kotrschal, K., 2002. Observational learning and the raiding of food caches in ravens, Corvus corax: is it ‘tactical’ deception? *Animal Behaviour*, [online] 64(2), pp.185–195. https://doi.org/10.1006/anbe.2002.3056.

Bugnyar, T. and Kotrschal, K., 2004. Leading a conspecific away from food in ravens *(Corvus corax)*? *Animal Cognition*, [online] 7(2), pp.69–76. https://doi.org/10.1007/s10071-003-0189-4.

Bugnyar, T., Reber, S.A. and Buckner, C., 2016. Ravens attribute visual access to unseen competitors. *Nature Communications*, [online] 7(1), p.10506. https://doi.org/10.1038/ncomms10506.

Bugnyar, T., Schwab, C., Schloegl, C., Kotrschal, K. and Heinrich, B., 2007. Ravens Judge Competitors through Experience with Play Caching. *Current Biology*, [online] 17(20), pp.1804–1808. https://doi.org/10.1016/j.cub.2007.09.048.

Bugnyar, T., Stöwe, M. and Heinrich, B., 2007. The ontogeny of caching in ravens, *Corvus corax*. *Animal Behaviour*, [online] 74(4), pp.757–767. https://doi.org/10.1016/j.anbehav.2006.08.019.

Burghardt, G. M. (2005). The genesis of animal play: Testing the limits. MIT press. https://mitpress.mit.edu/9780262524698/the-genesis-of-animal-play/

Carrascal, L. M., & Moreno, E. U. L. A. L. I. A. (1993). Food caching versus immediate consumption in the nuthatch: the effect of social context. ARDEA-WAGENINGEN*-*, 81, 135–135.

Cibulski, L., Wascher, C.A.F., Weiß, B.M. and Kotrschal, K., 2014. Familiarity with the experimenter influences the performance of Common ravens *(Corvus corax)* and Carrion crows *(Corvus corone corone)* in cognitive tasks. *Behavioural Processes*, [online] 103, pp.129–137. https://doi.org/10.1016/j.beproc.2013.11.013.

Clayton, N. and Emery, N., 2007. The social life of corvids. Current Biology, 17(16). Cramp, S. and Perrins, C., 1994. The birds of the Western Palearctic. Oxford: Oxford University Press.

Dally, J.M., Clayton, N.S. and Emery, N.J., 2006. The behaviour and evolution of cache protection and pilferage. *Animal Behaviour*, [online] 72(1), pp.13–23. https://doi.org/10.1016/j.anbehav.2005.08.020.

De Kort, S.R. and Clayton, N.S., 2006. An evolutionary perspective on caching by corvids. *Proceedings of the Royal Society B: Biological Sciences*, [online] 273(1585), pp.417–423. https://doi.org/10.1098/rspb.2005.3350.

De Kort, S.R., Emery, N.J. and Clayton, N.S., 2006. Food sharing in jackdaws, *Corvus monedula*: what, why and with whom? *Animal Behaviour*, [online] 72(2), pp.297–304. https://doi.org/10.1016/j.anbehav.2005.10.016.

Dukas, R. and Kamil, A.C., 2000. The cost of limited attention in blue jays. *Behavioral Ecology*, [online] 11(5), pp.502–506. https://doi.org/10.1093/beheco/11.5.502.

Emery, N.J., 2006. Cognitive ornithology: the evolution of avian intelligence. *Philosophical Transactions of the Royal Society B: Biological Sciences*, [online] 361(1465), pp.23–43. https://doi.org/10.1098/rstb.2005.1736.

Emery, N.J. and Clayton, N.S., 2002. Erratum: Effects of experience and social context on prospective caching strategies by scrub jays. *Nature*, [online] 416(6878), pp.349–349. https://doi.org/10.1038/416349a.

Ficken, M.S., 1977. Avian Play. *The Auk*, [online] 94(3), pp.573–582. https://doi.org/10.1093/auk/94.3.573.

Fite, K.V. and Rosenfield-Wessels, S., 1975. A Comparative Study of Deep Avian Foveas. *Brain, Behavior and Evolution*, [online] 12(1–2), pp.97–115. https://doi.org/10.1159/000124142.

Fransson, T., Kolehmainen, T., Kroon, C., Jansson, L. and Wenninger, T., 2010. EURING list of longevity records for European birds. http://www.euring.org/data_and_codes/longevity-voous.htm. [online] Available at: <https://cir.nii.ac.jp/crid/1572824499157708928> [Accessed 14 March 2023].

Fraser, O.N. and Bugnyar, T., 2010. The quality of social relationships in ravens. *Animal Behaviour*, [online] 79(4), pp.927–933. https://doi.org/10.1016/j.anbehav.2010.01.008.

Gajdon, G.K., Hungerbuhler, N. and Stauffacher, M., 2001. Social Influence on Early Foraging of Domestic Chicks *(Gallus gallus)* in a Near-to-Nature Procedure. *Ethology*, [online] 107(10), pp.913–937. https://doi.org/10.1046/j.1439-0310.2001.00719.x.

Gallego-Abenza, M., Boucherie, P.H. and Bugnyar, T., 2022. Early social environment affects attention to social cues in juvenile common ravens, *Corvus corax*. *Royal Society Open Science*, [online] 9(6), p.220132. https://doi.org/10.1098/rsos.220132.

Grampp, M., Sueur, C., van de Waal, E. and Botting, J., 2019. Social attention biases in juvenile wild vervet monkeys: implications for socialisation and social learning processes. *Primates*, [online] 60(3), pp.261–275. https://doi.org/10.1007/s10329-019-00721-4.

Hartig, F. and Lohse, L., 2022. DHARMa: Residual Diagnostics for Hierarchical (Multi-Level / Mixed) Regression Models. [online] Available at: <https://cran.r-project.org/web/packages/DHARMa/index.html>.

Heinrich, B. and Pepper, J.W., 1998. Influence of competitors on caching behaviour in the common raven, *Corvus corax*. *Animal Behaviour*, [online] 56(5), pp.1083–1090. https://doi.org/10.1006/anbe.1998.0906.

Heinroth, O. and Heinroth, M., 1926. Die Vögel Mitteleuropas [Birds of Central Europe]. Frankfurt: Verlag Harri Deutsch.

Heyes, C. M., & Galef Jr, B. G. (Eds.). (1996). Social learning in animals: the roots of culture. Elsevier. https://www.sciencedirect.com/book/9780122739651/social-learning-in-animals

Horn, L., Bugnyar, T., Griesser, M., Hengl, M., Izawa, E. I., Oortwijn, T., … & Massen, J. J. (2020). Sex-specific effects of cooperative breeding and colonial nesting on prosociality in corvids. Elife, 9, e58139. DOI: https://doi.org/10.7554/eLife.58139

Jacobs, I.F., Osvath, M., Osvath, H., Mioduszewska, B., Von Bayern, A.M.P. and Kacelnik, A., 2014. Object caching in corvids: Incidence and significance. *Behavioural Processes*, [online] 102, pp.25–32. https://doi.org/10.1016/j.beproc.2013.12.003.

Kano, F. and Call, J., 2017. Great Ape Social Attention. In: S. Watanabe, M.A. Hofman and T. Shimizu, eds. Evolution of the Brain, Cognition, and Emotion in Vertebrates, Brain Science. [online] Tokyo: Springer Japan. pp.187–206. https://doi.org/10.1007/978-4-431-56559-8_9.

Kano, F., Shepherd, S.V., Hirata, S. and Call, J., 2018. Primate social attention: Species differences and effects of individual experience in humans, great apes, and macaques. *PLOS ONE*, [online] 13(2), p.e0193283. https://doi.org/10.1371/journal.pone.0193283.

Kaplan, G., 2020. Play behaviour, not tool using, relates to brain mass in a sample of birds. *Scientific Reports*, [online] 10(1), p.20437. https://doi.org/10.1038/s41598-020-76572-7.

Klein, J.T., Shepherd, S.V. and Platt, M.L., 2009. Social Attention and the Brain. *Current Biology*, [online] 19(20), pp.R958–R962. https://doi.org/10.1016/j.cub.2009.08.010.

Kuczaj, S.A. and Eskelinen, H.C., 2014. Why do Dolphins Play? *Animal Behavior and Cognition*, [online] 2(2), p.113. https://doi.org/10.12966/abc.05.03.2014.

Kulahci, I.G., Rubenstein, D.I., Bugnyar, T., Hoppitt, W., Mikus, N. and Schwab, C., 2016. Social networks predict selective observation and information spread in ravens. *Royal Society Open Science*, [online] 3(7), p.160256. https://doi.org/10.1098/rsos.160256.

Lahti, K., Koivula, K., Rytkönen, S., Mustonen, T., Welling, P., Pravosudov, V.V. and Orell, M., 1998. Social influences on food caching in willow tits: a field experiment. *Behavioral Ecology*, [online] 9(2), pp.122–129. https://doi.org/10.1093/beheco/9.2.122.

Laland, K. N. (2004). Social learning strategies. Animal Learning & Behavior, 32, 4–14.

Lenth, R.V., Bolker, B., Buerkner, P., Giné-Vázquez, I., Herve, M., Jung, M., Love, J., Miguez, F., Riebl, H. and Singmann, H., 2023. emmeans: Estimated Marginal Means, aka Least-Squares Means. [online] Available at: <https://cran.rproject.org/web/packages/emmeans/index.html>.

Lima, S.L. and Bednekoff, P.A., 1999. Back to the basics of antipredatory vigilance: can nonvigilant animals detect attack? *Animal Behaviour*, [online] 58(3), pp.537–543. https://doi.org/10.1006/anbe.1999.1182.

Loretto, M.-C., Fraser, O.N. and Bugnyar, T., 2012. Ontogeny of Social Relations and Coalition Formation in Common Ravens *(Corvus corax). International Journal of Comparative Psychology*, [online] 25(3). https://doi.org/10.46867/IJCP.2012.25.03.05.

Loretto, M.-C., Schuster, R., Itty, C., Marchand, P., Genero, F. and Bugnyar, T., 2017. Fission-fusion dynamics over large distances in raven non-breeders. *Scientific Reports*, [online] 7(1), p.380. https://doi.org/10.1038/s41598-017-00404-4.

Marzluff, J.M. and Balda, R.P., 2010. The Pinyon Jay: Behavioral Ecology of a Colonial and Cooperative Corvid. AC Black.

Miller, R., Bugnyar, T., Pölzl, K. and Schwab, C., 2015. Differences in exploration behaviour in common ravens and carrion crows during development and across social context. *Behavioral Ecology and Sociobiology*, [online] 69(7), pp.1209–1220. https://doi.org/10.1007/s00265-015-1935-8.

Miller, R., Lambert, M.L., Frohnwieser, A., Brecht, K.F., Bugnyar, T., Crampton, I., Garcia-Pelegrin, E., Gould, K., Greggor, A.L., Izawa, E.-I., Kelly, D.M., Li, Z., Luo, Y., Luong, L.B., Massen, J.J.M., Nieder, A., Reber, S.A., Schiestl, M., Seguchi, A., Sepehri, P., Stevens, J.R., Taylor, A.H., Wang, L., Wolff, L.M., Zhang, Y. and Clayton, N.S., 2022. Socio-ecological correlates of neophobia in corvids. *Current Biology*, [online] 32(1), pp.74–85.e4. https://doi.org/10.1016/j.cub.2021.10.045.

Miller, R., Laskowski, K.L., Schiestl, M., Bugnyar, T. and Schwab, C., 2016. Socially Driven Consistent Behavioural Differences during Development in Common Ravens and Carrion Crows. *PLOS ONE*, [online] 11(2), p.e0148822. https://doi.org/10.1371/journal.pone.0148822.

Moscovice, L.R. and Snowdon, C.T., 2006. The role of social context and individual experience in novel task acquisition in cottontop tamarins, *Saguinus oedipus*. *Animal Behaviour*, [online] 71(4), pp.933–943. https://doi.org/10.1016/j.anbehav.2005.09.007.

Naples, V. and Rothschild, B., 2015. Play Behaviour in Primates. Primatology, [online] 4. Available at: <https://www.omicsgroup.org/journals/play-behavior-in-primates-2167-6801-1000e132.pdf> [Accessed 20 June 2023].

Nunes, S., Muecke, E.-M., Sanchez, Z., Hoffmeier, R.R. and Lancaster, L.T., 2004. Play behavior and motor development in juvenile Belding’s ground squirrels *(Spermophilus beldingi)*. *Behavioral Ecology and Sociobiology*, [online] 56(2), pp.97–105. https://doi.org/10.1007/s00265-004-0765-x.

O’Hara, M. and Auersperg, A.M., 2017. Object play in parrots and corvids. *Current Opinion in Behavioral Sciences*, [online] 16, pp.119–125. https://doi.org/10.1016/j.cobeha.2017.05.008.

Oliva, A., Torralba, A., Castelhano, M.S. and Henderson, J.M., 2003. Top-down control of visual attention in object detection. In: Proceedings 2003 International Conference on Image Processing (Cat. No.03CH37429). [online] International Conference on Image Processing. Barcelona, Spain: IEEE. p.I-253–6. https://doi.org/10.1109/ICIP.2003.1246946.

Palagi, E., 2018. Not just for fun! Social play as a springboard for adult social competence in human and non-human primates. *Behavioral Ecology and Sociobiology*, [online] 72(6), p.90. https://doi.org/10.1007/s00265-018-2506-6.

Palagi, E., Cordoni, G. and Borgognini Tarli, S.M., 2004. Immediate and Delayed Benefits of Play Behaviour: New Evidence from Chimpanzees (*Pan troglodytes*). *Ethology*, [online] 110(12), pp.949–962. https://doi.org/10.1111/j.1439-0310.2004.01035.x.

Pellegrini, A.D. and Smith, P.K., 2005. The Nature of Play: Great Apes and Humans. Guilford Press.

Poirier, F.E. and Smith, E.O., 1974. Socializing Functions of Primate Play. *American Zoologist*, [online] 14(1), pp.275–287. https://doi.org/10.1093/icb/14.1.275.

Ramsey, J. K. and McGrew, W. C., 2005. Object Play in Great Apes: Studies in Nature and Captivity. In A. D. Pellegrini and P. K. Smith, ed. 2005. The nature of play: Great apes and humans. New York: The Guilford Press, pp.89–112.

Range, F., Horn, L., Bugnyar, T., Gajdon, G.K. and Huber, L., 2009. Social attention in keas, dogs, and human children. *Animal Cognition*, [online] 12(1), pp.181–192. https://doi.org/10.1007/s10071-008-0181-0.

Ratcliff, D., 1997. The raven. San Diego: Academic Press Inc.

R Core Team, 2023. R: A Language and Environment for Statistical Computing, R Foundation for Statistical Computing, [online] Vienna, Austria. Available at: <https://www.R-project.org/>.

Richardson, D.C. and Globel, M.S., 2015. The Handbook of Attention. MIT Press.

RSPB, 2023a. Carrion Crow Facts | Corvus Corone. [online] The RSPB. Available at: <https://www.rspb.org.uk/birds-and-wildlife/wildlife-guides/bird-a-z/carrion-crow/> [Accessed 2 July 2023].

RSPB, 2023b. Common Raven Bird Facts | Corvus Corax. [online] The RSPB. Available at: <https://www.rspb.org.uk/birds-and-wildlife/wildlife-guides/bird-a-z/raven/> [Accessed 2 July 2023].

Scheid, C., Range, F. and Bugnyar, T., 2007. When, what, and whom to watch? Quantifying attention in ravens *(Corvus corax)* and jackdaws *(Corvus monedula). Journal of Comparative Psychology*, [online] 121(4), pp.380–386. https://doi.org/10.1037/0735-7036.121.4.380.

Scheid, C., Schmidt, J. and Noë, R., 2008. Distinct patterns of food offering and co-feeding in rooks. *Animal Behaviour*, [online] 76(5), pp.1701–1707. https://doi.org/10.1016/j.anbehav.2008.07.023.

Schino, G. and Sciarretta, M., 2016. Patterns of Social Attention in Mandrills, *Mandrillus sphinx*. *International Journal of Primatology*, [online] 37(6), pp.752–761. https://doi.org/10.1007/s10764-016-9936-7.

Schloegl, C., Kotrschal, K., & Bugnyar, T. (2007). Gaze following in common ravens, Corvus corax: ontogeny and habituation. Animal Behaviour, 74(4), 769–778. https://doi.org/10.1016/j.anbehav.2006.08.017

Schwab, C., Bugnyar, T. and Kotrschal, K., 2008. Preferential learning from non-affiliated individuals in jackdaws *(Corvus monedula). Behavioural Processes*, [online] 79(3), pp.148–155. https://doi.org/10.1016/j.beproc.2008.07.002.

Schwab, C., Bugnyar, T., Schloegl, C. and Kotrschal, K., 2008. Enhanced social learning between siblings in common ravens, Corvus corax. *Animal Behaviour*, [online] 75(2), pp.501–508. https://doi.org/10.1016/j.anbehav.2007.06.006.

Silk, M.J., Croft, D.P., Tregenza, T. and Bearhop, S., 2014. The importance of fission-fusion social group dynamics in birds. *Ibis*, [online] 156(4), pp.701–715. https://doi.org/10.1111/ibi.12191.

Smith, P.K., 1982. Does play matter? Functional and evolutionary aspects of animal and human play. *Behavioral and Brain Sciences*, [online] 5(1), pp.139–155. https://doi.org/10.1017/S0140525X0001092X.

Stöwe, M., Bugnyar, T., Heinrich, B. and Kotrschal, K., 2006. Effects of Group Size on Approach to Novel Objects in Ravens *(Corvus corax)*. *Ethology*, [online] 112(11), pp.1079– 1088. https://doi.org/10.1111/j.1439-0310.2006.01273.x.

Taborsky, B. and Oliveira, R.F., 2012. Social competence: an evolutionary approach. *Trends in Ecology & Evolution*, [online] 27(12), pp.679–688. https://doi.org/10.1016/j.tree.2012.09.003.

Uhl, F., Ringler, M., Miller, R., Deventer, S.A., Bugnyar, T. and Schwab, C., 2019. Counting crows: population structure and group size variation in an urban population of crows.*Behavioral Ecology*, [online] 30(1), pp.57–67. https://doi.org/10.1093/beheco/ary157.

Vander Wall, S. B., & Smith, K. G. (1987). Cache-protecting behavior of food-hoarding animals. Foraging behavior, 611–644.

von Bayern, A. M., de Kort, S. R., Clayton, N. S., & Emery, N. J. (2007). The role of food-and object-sharing in the development of social bonds in juvenile jackdaws (Corvus monedula). Behaviour, 711–733. https://brill.com/view/journals/beh/144/6/article-p711_6.xml?language=en

Wascher, C.A.F., Valdez, J.W., Cebrián, C.N., Baglione, V. and Canestrari, D., 2014. Social factors modulating attention patterns in carrion crows. *Behaviour*, [online] 151(5), pp.555–572. https://doi.org/10.1163/1568539X-00003148.

Whiten, A., Goodall, J., McGrew, W. C., Nishida, T., Reynolds, V., Sugiyama, Y., … & Boesch, C. (1999). Cultures in chimpanzees. Nature, 399(6737), 682–685. https://doi.org/10.1038/21415

Whiten, A., & Van Schaik, C. P. (2007). The evolution of animal ‘cultures’ and social intelligence. Philosophical Transactions of the Royal Society B: Biological Sciences, 362(1480), 603–620. doi: 10.1098/rstb.2006.1998

