## Supplementary Materials for "Social attention across development in common ravens and carrion crows"

**Supplementary Table 1: Subject Information.** Ns = no-sibling so not tested as focal in round 1 and 2. / = died so not tested in round 3. Affiliate = most affiliative interactions; non-affiliate = no or least affiliative interactions

| **ID** | **Species** | **Sex** | **Tested in round** | **Sibling (R1&2)** | **Affiliate (R3)** | **Non-sibling (R1&2)** | **Non-affiliate (R3)** | **Heterospecific** |
| --- | --- | --- | --- | --- | --- | --- | --- | --- |
| Adele | Common raven | F | 1-3 | Laggie | Laggie | Horst | Louise | Saul |
| Laggie | Common raven | M | 1-3 | Tom | Tom | George | Horst | Saul |
| Tom | Common raven | M | 1-3 | Laggie | Adele | George | Horst | Signore |
| George | Common raven | M | 1-3 | Horst | Nobel | Laggie | Adele | Saul |
| Horst | Common raven | M | 1-3 | George | Louise | Paul | Rufous | Corbie |
| Louise | Common raven | F | 1-3 | Nobel | George | Adele | Rufous | Suki |
| Nobel | Common raven | F | 1-3 | Louise | Horst | Adele | Paul | Daisy |
| Paul | Common raven | M | 1-3 | Max | Max | Laggie | Louise | Corbie |
| Rufous | Common raven | M | 1-3 | ns | Adele | ns | Nobel | Corbie |
| Corbie | Carrion crow | M | 1-3 | ns | Peppi | ns | Saul | Rufous |
| Saul | Carrion crow | M | 1-3 | Signore | Rainer | Corbie | Daisy | Horst |
| Signore | Carrion crow | M | 1-3 | Saul | Rainer | Corbie | Daisy | George |
| Daisy | Carrion crow | F | 1-3 | Suki | Corbie | Peppi | Saul | Louise |
| Emily | Carrion crow | F | 1-2 | Daisy | / | Peppi | / | Nobel |
| Suki | Carrion crow | F | 1-3 | Emily | Signore | Rainer | Corbie | Louise |
| Juno | Carrion crow | F | 2-3 | Lilith | Rainer | Suki | Daisy | Nobel |
| Lilith | Carrion crow | M | 2 | Juna | / | Emily | / | Nobel |
| Peppi | Carrion crow | F | 1-2 | Rainer | Saul | Suki | Rainer | Louise |
| Rainer | Carrion crow | F | 1-3 | Peppi | Corbie | Daisy | Suki | Adele |

**Supplementary Table 2: GLMM model output on factors affecting 3 behavioural measures (item manipulation, caching, head & body out of sight).** P<0.05 highlighted in bold

| **GLMM/ Behaviour measure** | **Effect** | **Estimate** | **z** | **p** |
| --- | --- | --- | --- | --- |
| Item Manipulation | Species1 | 0.057 | 0.278 | 0.781 |
|  | Context2 | 0.172 | 1.222 | 0.222 |
|  | Context3 | 0.231 | 1.615 | 0.106 |
|  | Context4 | 0.211 | 1.545 | 0.122 |
|  | Item type1 | -0.464 | -6.516 | **<0.0001** |
|  | Round2 | 0.135 | 1.141 | 0.254 |
|  | Round3 | -0.069 | -0.561 | 0.575 |
|  | Species1*Round2 | 0.468 | 2.642 | **0.008** |
|  | Species1*Round3 | 0.320 | 1.695 | 0.090 |
|  | Species1*Context2 | -0.307 | -1.552 | 0.121 |
|  | Species1*Context3 | -0.466 | -2.316 | **0.021** |
|  | Species1*Context4 | -0.232 | -1.203 | 0.229 |
| Caching | Species1 | -0.955 | -2.706 | **0.007** |
|  | Context2 | 0.322 | 1.718 | 0.086 |
|  | Context3 | 0.152 | 0.811 | 0.417 |
|  | Context4 | 0.029 | 0.152 | 0.880 |
|  | Item type1 | 0.202 | 1.910 | 0.056 |
|  | Round2 | 0.309 | 1.621 | 0.105 |
|  | Round3 | 1.105 | 6.545 | **<0.0001** |
|  | Species1*Round2 | 1.305 | 3.884 | **0.0001** |
|  | Species1*Round3 | 0.628 | 1.938 | 0.053 |
|  | Species1*Context2 | -0.503 | -1.776 | 0.076 |
|  | Species1*Context3 | -0.214 | -0.775 | 0.438 |
|  | Species1*Context4 | -0.291 | -1.033 | 0.302 |
| Head and Body  out of sight | Species1 | -0.532 | -1.536 | 0.125 |
|  | Context2 | 0.745 | 2.871 | **0.004** |
|  | Context3 | 0.719 | 2.727 | **0.006** |
|  | Context4 | 0.647 | 2.501 | **0.012** |
|  | Item type1 | 1.001 | 7.468 | **<0.0001** |
|  | Round2 | 0.715 | 3.155 | **0.002** |
|  | Round3 | 0.092 | 0.419 | 0.676 |
|  | Species1*Round2 | 0.856 | 2.416 | **0.016** |
|  | Species1*Round3 | 0.708 | 2.081 | **0.037** |
|  | Species1*Context2 | -0.060 | -0.166 | 0.868 |
|  | Species1*Context3 | -0.081 | -0.223 | 0.823 |
|  | Species1*Context4 | -0.203 | -0.578 | 0.564 |

**Supplementary Table 3. Tukey contrasts pairwise comparisons with main effects.**

P<0.05 highlighted in bold. 3 behavioural measures: item manipulation, caching and head & body out of sight.

| **Behaviour** | **Fixed effect** | **Comparison** | **z** | **p** |
| --- | --- | --- | --- | --- |
| Manipulation | Species | 0-1 | -0.609 | 0.543 |
| Caching | Species | 0-1 | 3.301 | **0.001** |
| HeadBodyOOS | Species | 0-1 | 0.730 | 0.466 |
| Manipulation | Item type | 0-1 | 6.516 | **<0.0001** |
| Caching | Item type | 0-1 | -1.910 | *0.0561* |
| HeadBodyOOS | Item type | 0-1 | -7.468 | **<0.0001** |
| Manipulation | Round | 1-2 | -4.185 | **0.0001** |
| Manipulation | Round | 1-3 | -0.963 | 0.6002 |
| Manipulation | Round | 2-3 | 3.2229 | **0.0036** |
| Caching | Round | 1-2 | -5.816 | **<0.0001** |
| Caching | Round | 1-3 | -8.706 | **<0.0001** |
| Caching | Round | 2-3 | -3.980 | **0.0002** |
| HeadBodyOOS | Round | 1-2 | -7.258 | **<0.0001** |
| HeadBodyOOS | Round | 1-3 | -2.613 | **0.0244** |
| HeadBodyOOS | Round | 2-3 | 4.649 | **<0.0001** |
| Manipulation | Social context | 1-2 | -0.180 | 0.998 |
| Manipulation | Social context | 1-3 | 0.016 | >0.999 |
| Manipulation | Social context | 1-4 | -0.984 | 0.759 |
| Manipulation | Social context | 2-3 | 0.194 | 0.997 |
| Manipulation | Social context | 2-4 | -0.801 | 0.854 |
| Manipulation | Social context | 3-4 | -0.985 | 0.758 |
| Caching | Social context | 1-2 | -.500 | 0.959 |
| Caching | Social context | 1-3 | -0.325 | 0.988 |
| Caching | Social context | 1-4 | 0.823 | 0.843 |
| Caching | Social context | 2-3 | 0.178 | 0.998 |
| Caching | Social context | 2-4 | 1.279 | 0.576 |
| Caching | Social context | 3-4 | 1.116 | 0.679 |
| HeadBodyOOS | Social context | 1-2 | -3.771 | **0.0009** |
| HeadBodyOOS | Social context | 1-3 | -3.662 | **0.0014** |
| HeadBodyOOS | Social context | 1-4 | -3.012 | **0.0138** |
| HeadBodyOOS | Social context | 2-3 | 0.213 | 0.997 |
| HeadBodyOOS | Social context | 2-4 | 1.001 | 0.749 |
| HeadBodyOOS | Social context | 3-4 | 0.789 | 0.859 |

**Supplementary Table 4. Tukey contrasts pairwise comparisons with interaction effects.** R = raven; C = crow. p<0.05 highlighted in bold.

| **Behaviour** | **Interaction** | **Comparison** | | **z** | **p** |
| --- | --- | --- | --- | --- | --- |
| Manipulation | Species*Context | R.1 | C.1 | -1.920 | 0.537 |
| Manipulation | Species*Context | R.1 | R.2 | -1.222 | 0.926 |
| Manipulation | Species*Context | R.1 | C.2 | -1.103 | 0.956 |
| Manipulation | Species*Context | R.1 | R.3 | -1.615 | 0.742 |
| Manipulation | Species*Context | R.1 | C.3 | -0.506 | 1.000 |
| Manipulation | Species*Context | R.1 | R.4 | -1.545 | 0.783 |
| Manipulation | Species*Context | R.1 | C.4 | -1.818 | 0.608 |
| Manipulation | Species*Context | C.1 | R.2 | 0.900 | 0.986 |
| Manipulation | Species*Context | C.1 | C.2 | 0.969 | 0.979 |
| Manipulation | Species*Context | C.1 | R.3 | 0.532 | 1.000 |
| Manipulation | Species*Context | C.1 | C.3 | 1.660 | 0.713 |
| Manipulation | Species*Context | C.1 | R.4 | 0.671 | 0.998 |
| Manipulation | Species*Context | C.1 | C.4 | 0.154 | 1.000 |
| Manipulation | Species*Context | R.2 | C.2 | -0.076 | 1.000 |
| Manipulation | Species*Context | R.2 | R.3 | -0.424 | 1.000 |
| Manipulation | Species*Context | R.2 | C.3 | 0.514 | 1.000 |
| Manipulation | Species*Context | R.2 | R.4 | -0.294 | 1.000 |
| Manipulation | Species*Context | R.2 | C.4 | -0.784 | 0.994 |
| Manipulation | Species*Context | C.2 | R.3 | -0.282 | 1.000 |
| Manipulation | Species*Context | C.2 | C.3 | 0.689 | 0.997 |
| Manipulation | Species*Context | C.2 | R.4 | -0.167 | 1.000 |
| Manipulation | Species*Context | C.2 | C.4 | -0.832 | 0.991 |
| Manipulation | Species*Context | R.3 | C.3 | 0.862 | 0.989 |
| Manipulation | Species*Context | R.3 | R.4 | 0.146 | 1.000 |
| Manipulation | Species*Context | R.3 | C.4 | -0.412 | 1.000 |
| Manipulation | Species*Context | C.3 | R.4 | -0.765 | 0.995 |
| Manipulation | Species*Context | C.3 | C.4 | -1.524 | 0.795 |
| Manipulation | Species*Context | R.4 | C.4 | -0.550 | 0.995 |
| Manipulation | Species*Round | R.1 | R.2 | -1.141 | 0.864 |
| Manipulation | Species*Round | R.1 | R.3 | 0.561 | 0.994 |
| Manipulation | Species*Round | R.2 | R.3 | 1.655 | 0.562 |
| Manipulation | Species*Round | C.1 | C.2 | -4.584 | **0.0001** |
| Manipulation | Species*Round | C.1 | C.3 | -1.761 | 0.491 |
| Manipulation | Species*Round | C.2 | C.3 | 2.909 | **0.042** |
| Manipulation | Species*Round | R.1 | C.1 | 1.196 | 0.839 |
| Manipulation | Species*Round | R.1 | C.2 | -2.888 | **0.045** |
| Manipulation | Species*Round | R.1 | C.3 | -0.371 | 0.999 |
| Manipulation | Species*Round | R.2 | C.1 | 2.035 | 0.322 |
| Manipulation | Species*Round | R.2 | C.2 | -1.940 | 0.377 |
| Manipulation | Species*Round | R.2 | C.3 | 0.515 | 0.996 |
| Manipulation | Species*Round | R.3 | C.1 | 0.758 | 0.974 |
| Manipulation | Species*Round | R.3 | C.2 | -3.305 | **0.012** |
| Manipulation | Species*Round | R.3 | C.3 | -0.812 | 0.965 |
| Caching | Species*Context | R.1 | C.1 | 1.317 | 0.892 |
| Caching | Species*Context | R.1 | R.2 | -1.718 | 0.676 |
| Caching | Species*Context | R.1 | C.2 | 1.976 | 0.498 |
| Caching | Species*Context | R.1 | R.3 | -0.811 | 0.993 |
| Caching | Species*Context | R.1 | C.3 | 1.538 | 0.787 |
| Caching | Species*Context | R.1 | R.4 | -0.152 | 1.000 |
| Caching | Species*Context | R.1 | C.4 | 2.326 | 0.279 |
| Caching | Species*Context | C.1 | R.2 | -2.688 | 0.126 |
| Caching | Species*Context | C.1 | C.2 | 0.858 | 0.990 |
| Caching | Species*Context | C.1 | R.3 | -1.965 | 0.506 |
| Caching | Species*Context | C.1 | C.3 | 0.307 | 1.000 |
| Caching | Species*Context | C.1 | R.4 | -1.421 | 0.848 |
| Caching | Species*Context | C.1 | C.4 | 1.260 | 0.913 |
| Caching | Species*Context | R.2 | C.2 | 3.265 | **0.024** |
| Caching | Species*Context | R.2 | R.3 | 0.914 | 0.985 |
| Caching | Species*Context | R.2 | C.3 | 2.869 | 0.079 |
| Caching | Species*Context | R.2 | R.4 | 1.555 | 0.777 |
| Caching | Species*Context | R.2 | C.4 | 3.656 | **0.006** |
| Caching | Species*Context | C.2 | R.3 | -2.587 | 0.160 |
| Caching | Species*Context | C.2 | C.3 | -0.544 | 0.999 |
| Caching | Species*Context | C.2 | R.4 | -2.064 | 0.439 |
| Caching | Species*Context | C.2 | C.4 | 0.361 | 1.000 |
| Caching | Species*Context | R.3 | C.3 | 2.168 | 0.371 |
| Caching | Species*Context | R.3 | R.4 | 0.643 | 0.998 |
| Caching | Species*Context | R.3 | C.4 | 2.952 | 0.063 |
| Caching | Species*Context | C.3 | R.4 | -1.635 | 0.729 |
| Caching | Species*Context | C.3 | C.4 | 0.925 | 0.984 |
| Caching | Species*Context | R.4 | C.4 | 2.423 | 0.230 |
| Caching | Species*Round | R.1 | R.2 | -1.621 | 0.585 |
| Caching | Species*Round | R.1 | R.3 | -6.545 | **<0.0001** |
| Caching | Species*Round | R.2 | R.3 | -4.785 | **<0.0001** |
| Caching | Species*Round | C.1 | C.2 | -5.902 | **<0.0001** |
| Caching | Species*Round | C.1 | C.3 | -6.237 | **<0.0001** |
| Caching | Species*Round | C.2 | C.3 | -0.731 | 0.978 |
| Caching | Species*Round | R.1 | C.1 | 3.908 | **0.001** |
| Caching | Species*Round | R.1 | C.2 | -1.920 | 0.390 |
| Caching | Species*Round | R.1 | C.3 | -2.420 | 0.149 |
| Caching | Species*Round | R.2 | C.1 | 4.942 | **<0.0001** |
| Caching | Species*Round | R.2 | C.2 | -0.459 | 0.998 |
| Caching | Species*Round | R.2 | C.3 | -1.002 | 0.918 |
| Caching | Species*Round | R.3 | C.1 | 7.864 | **<0.0001** |
| Caching | Species*Round | R.3 | C.2 | 3.746 | **0.003** |
| Caching | Species*Round | R.3 | C.3 | 2.984 | **0.034** |
| HeadBodyOOS | Species*Context | R.1 | C.1 | 0.04 | 1.000 |
| HeadBodyOOS | Species*Context | R.1 | R.2 | -2.871 | 0.079 |
| HeadBodyOOS | Species*Context | R.1 | C.2 | -2.509 | 0.192 |
| HeadBodyOOS | Species*Context | R.1 | R.3 | -2.727 | 0.114 |
| HeadBodyOOS | Species*Context | R.1 | C.3 | -2.363 | 0.260 |
| HeadBodyOOS | Species*Context | R.1 | R.4 | -2.501 | 0.195 |
| HeadBodyOOS | Species*Context | R.1 | C.4 | -1.685 | 0.697 |
| HeadBodyOOS | Species*Context | C.1 | R.2 | -2.951 | 0.063 |
| HeadBodyOOS | Species*Context | C.1 | C.2 | -2.604 | 0.154 |
| HeadBodyOOS | Species*Context | C.1 | R.3 | -2.808 | 0.093 |
| HeadBodyOOS | Species*Context | C.1 | C.3 | -2.489 | 0.199 |
| HeadBodyOOS | Species*Context | C.1 | R.4 | -2.575 | 0.165 |
| HeadBodyOOS | Species*Context | C.1 | C.4 | -1.808 | 0.615 |
| HeadBodyOOS | Species*Context | R.2 | C.2 | 0.283 | 1.000 |
| HeadBodyOOS | Species*Context | R.2 | R.3 | 0.103 | 1.000 |
| HeadBodyOOS | Species*Context | R.2 | C.3 | 0.481 | 1.000 |
| HeadBodyOOS | Species*Context | R.2 | R.4 | 0.412 | 1.000 |
| HeadBodyOOS | Species*Context | R.2 | C.4 | 1.288 | 0.904 |
| HeadBodyOOS | Species*Context | C.2 | R.3 | -0.169 | 1.000 |
| HeadBodyOOS | Species*Context | C.2 | C.3 | 0.204 | 1.000 |
| HeadBodyOOS | Species*Context | C.2 | R.4 | 0.117 | 1.000 |
| HeadBodyOOS | Species*Context | C.2 | C.4 | 0.993 | 0.976 |
| HeadBodyOOS | Species*Context | R.3 | C.3 | 0.356 | 1.000 |
| HeadBodyOOS | Species*Context | R.3 | R.4 | 0.291 | 1.000 |
| HeadBodyOOS | Species*Context | R.3 | C.4 | 1.150 | 0.946 |
| HeadBodyOOS | Species*Context | C.3 | R.4 | -0.078 | 1.000 |
| HeadBodyOOS | Species*Context | C.3 | C.4 | 0.820 | 0.992 |
| HeadBodyOOS | Species*Context | R.4 | C.4 | 0.887 | 0.987 |
| HeadBodyOOS | Species*Round | R.1 | R.2 | -3.155 | **0.020** |
| HeadBodyOOS | Species*Round | R.1 | R.3 | -0.419 | 0.998 |
| HeadBodyOOS | Species*Round | R.2 | R.3 | 2.680 | 0.079 |
| HeadBodyOOS | Species*Round | C.1 | C.2 | -6.357 | **<0.0001** |
| HeadBodyOOS | Species*Round | C.1 | C.3 | -3.076 | **0.026** |
| HeadBodyOOS | Species*Round | C.2 | C.3 | 3.276 | **0.013** |
| HeadBodyOOS | Species*Round | R.1 | C.1 | 2.486 | 0.128 |
| HeadBodyOOS | Species*Round | R.1 | C.2 | -4.610 | **0.0001** |
| HeadBodyOOS | Species*Round | R.1 | C.3 | -0.784 | 0.970 |
| HeadBodyOOS | Species*Round | R.2 | C.1 | 5.299 | **<0.0001** |
| HeadBodyOOS | Species*Round | R.2 | C.2 | -0.995 | 0.920 |
| HeadBodyOOS | Species*Round | R.2 | C.3 | 2.330 | 0.182 |
| HeadBodyOOS | Species*Round | R.3 | C.1 | 2.813 | 0.055 |
| HeadBodyOOS | Species*Round | R.3 | C.2 | -4.198 | **0.0004** |
| HeadBodyOOS | Species*Round | R.3 | C.3 | -0.383 | 0.999 |
